# Understanding functional consequences of type 2 diabetes risk loci using the universal data integration and visualization R package CONQUER

**DOI:** 10.1101/2020.03.27.011627

**Authors:** Gerard A Bouland, Joline WJ Beulens, Joey Nap, Arno R van der Slik, Arnaud Zaldumbide, Leen M’t Hart, Roderick C Slieker

## Abstract

**Background:** Numerous large genome-wide association studies (GWASs) have been performed to understand the genetic factors of numerous traits, including type 2 diabetes. Many identified risk loci are located in non-coding and intergenic regions, which complicates the understanding how genes and their downstream pathways are influenced. An integrative data approach is required to understand the mechanism and consequences of identified risk loci.

**Results:** Here, we developed the R-package CONQUER. Data for SNPs of interest (build GRCh38/hg38) were acquired from static- and dynamic repositories, such as, GTExPortal, Epigenomics Project, 4D genome database and genome browsers such as ENSEMBL. CONQUER modularizes SNPs based on the underlying co-expression data and associates them with biological pathways in specific tissues. CONQUER was used to analyze 403 previously identified type 2 diabetes risk loci. In all tissues, the majority of SNPs (mean = 13.50, SD = 11.70) were linked to metabolism. A tissue-shared effect was found for four type 2 diabetes-associated SNPs (rs601945, rs1061810, rs13737, rs4932265) that were associated with differential expression of *HLA-DQA2, HSD17B12, MAN2C1* and *AP3S2* respectively. Seven SNPs were identified that influenced the expression of seven ribosomal proteins in multiple tissues. Finally, one SNP (rs601945) was found to influence multiple *HLA* genes in all twelve tissues investigated.

**Conclusion:** We present an universal R-package that aggregates and visualizes data in order to better understand functional consequences of GWAS loci. Using CONQUER, we showed that type 2 diabetes risk loci have many tissue-shared effects on multiple pathways including metabolism, the ribosome and HLA pathway.

## BACKGROUND

In the past decades, numerous genome-wide association studies (GWAS) have been performed to understand the genetic contribution of traits. While GWASs have provided valuable insight into putative mechanistic pathways, the way the identified risk loci exert their effect on traits remain largely unclear. For type 2 diabetes (T2D), several large meta-GWASs have been performed to understand the genetic drivers of T2D [1–3]. In general, GWAS associated loci are not limited to coding regions but are frequently found in intergenic regions [4]. As such, inferring how risk loci influence genes and their downstream pathways remains often unclear, especially for loci in non-coding regions. To increase the understanding of those variants, an integrative approach is required where the effects of variants are investigated at a multitude of molecular levels.

In recent years, the number of rich open source biological data sets and repositories has tremendously increased, including GTExPortal [5], Epigenomics Project [6], 4D genome database[7] and genome browsers such as ENSEMBL [8]. Extracting, combining and analyzing relevant biological information from these public datasets is complicated and time-consuming. Platforms that integrate such data exist [9, 10], but are often online, miss intuitive user experience or contain outdated data or genome builds. To provide researchers with a easy to use interface with the latest data to comprehend the effects of variants, we developed an R-package named CONQUER (‘COmprehend fuNctional conseQUencEs R’). Given a single SNP or multiple SNPs, CONQUER allows the user to efficiently extract relevant biological information from various repositories/databases and represents the information through insightful and interactive visualizations. Additionally, CONQUER links SNPs with biological pathways trough enrichment of the associated genes. Here, we use CONQUER to investigate the 403 risk loci associated with T2D in more detail. With CONQUER we identified T2D risk loci that influence the expression of genes in diabetes-relevant tissues.

## RESULTS

### CONQUER: an universal R-package for GWAS loci

CONQUER is a universal tool that retrieves and visualizes a multitude of public data associated with any SNP of interest. The package can be used both for single and multiple SNPs. In both cases, CONQUER collects data about a SNP from various public databases and stores the data locally per SNP in a file. Data is collected on multiple levels, including expression-, methylation, metabolomics- and protein QTLs, chromosomal interactions, histone modifications and GWAS catalogue (see methods). All tissues included in GTEx can be investigated with CONQUER. When multiple SNPs are investigated, CONQUER will find shared pathways across the SNPs investigated (see methods). Results are integrated in an interactive offline web interface for the analysis of multiple SNPs (https://github.com/roderickslieker/CONQUER). Altogether, CONQUER has two separate views 1) where in-depth analyses of single SNPs can performed and 2) where multiple SNPs and their aggregated consequences can be investigated and linked to biological pathways. Here, CONQUER was used to analyze 403 T2D-associated SNPs.

### Type 2 diabetes-associated eQTLs are tissue-shared

From the most recent meta-GWAS, 403 T2D-associated SNPs were obtained [1]. Out of those, 17 SNPs were associated with in total 23 unique pQTLs (22 trans, 1 cis). Of those, nine were associated with the immune system (REACTOME, *P*=0.01), including IL17RC, ICAM1, SAA1, ULBP1, C3, DSG1, CFI, IL18RAP, MBL2. Two trans pQTLs were involved in cholesterol metabolism, LPA and ANGPTL3. Of note, the LPA protein was the single *cis* signal and associated with rs474513 (*P*=8.27·10^−37^). Although this variant is an eQTL in 17 tissues for SLC22A2, it was an eQTLs for *LPA* in the liver (*P*=1.29·10^−5^). SLC22A2 encodes the organic cation transporter 2 gene (OCT2) which is involved in the uptake of the glucose-lowering drug metformin in the kidneys [11] and *LPA* encodes the lipoprotein A protein which is thought to be atherogenic [12].

Next, SNPs were investigated in gene expression data of tissues relevant in the etiology of diabetes (subcutaneous and visceral fat, sigmoid- and transverse colon, liver, skeletal muscle, pancreas, pituitary, terminal ileum of the small intestine, stomach, thyroid and whole blood) from healthy individuals. Of the included tissues, sample sizes range from N = 187 (terminal ileum) to N = 803 (skeletal muscle). Characteristics are shown in Table 1. The percentage males was relatively equal across tissues (63.1% - 72.1%, **Table 1**) with the majority middle-aged (50-69 years, **Table 1**). For the eQTL – eGene analysis, CONQUER retrieved 348 SNPs. Fifty-five SNPs were excluded because they were not a significant eQTL (33 SNPs) or the variant ID was not present (23 SNPs). Sample sizes are correlated with the number of significant eQTLs (R^2^=0.91). We take this into account when evaluating the results by applying a liberal threshold (P≤0.001) and by assessing the normalized effect sizes (NES) across tissues. Using the 348 SNPs, cis- and trans genes were calculated with the GTEx API. After applying a threshold (P ≤ 0.05), sets of co-expressed genes were determined after which all included genes (eGenes and co-expressed genes) were clustered. Out of the 348 SNPs, 214 SNPs were significant. This resulted in 6664 calculated eQTL - eGene pairs across tissues (**Fig.1a**).

**Table 1.** Characteristics of the individuals in the GTEX data.

| Tissue | N | Sex (male) | Age |  |  |  |  |  | 0 |  |
| --- | --- | --- | --- | --- | --- | --- | --- | --- | --- | --- |
|  |  |  | 20-29 | 30-39 | 40-49 | 50-59 | 60-69 | 70-79 |  |  |
| Adipose Subcutaneous | 663 | 445 (67,1 %) | 52 (7,8 %) | 58 (8,7 %) | 103 (15,5 %) | 213 (32,1 %) | 216 (32,6 %) | 21 (3,2 %) | 352 (53,1 %) | 21 (3,2 %) |
| Adipose Visceral Omentum | 541 | 371 (68,6 %) | 45 (8,3 %) | 44 (8,1 %) | 86 (15,9 %) | 184 (34 %) | 163 (30,1 %) | 19 (3,5 %) | 314 (58 %) | 16 (3,0 %) |
| Colon Sigmoid | 373 | 240 (64,3 %) | 36 (9,7 %) | 39 (10,5 %) | 54 (14,5 %) | 109 (29,2 %) | 123 (33 %) | 12 (3,2 %) | 236 (63,3 %) | 9 (2,4 %) |
| Colon Transverse | 406 | 259 (63,8 %) | 43 (10,6 %) | 47 (11,6 %) | 75 (18,5 %) | 136 (33,5 %) | 94 (23,2 %) | 11 (2,7 %) | 316 (77,8 %) | 4 (1,0 %) |
| Liver | 226 | 161 (71,2 %) | 7 (3,1 %) | 17 (7,5 %) | 35 (15,5 %) | 83 (36,7 %) | 79 (35 %) | 5 (2,2 %) | 86 (38,1 %) | 13 (5,8 %) |
| Muscle Skeletal | 803 | 543 (67,6 %) | 67 (8,3 %) | 65 (8,1 %) | 124 (15,4 %) | 255 (31,8 %) | 264 (32,9 %) | 28 (3,5 %) | 424 (52,8 %) | 31 (3,9 %) |
| Pancreas | 328 | 207 (63,1 %) | 29 (8,8 %) | 31 (9,5 %) | 66 (20,1 %) | 118 (36 %) | 79 (24,1 %) | 5 (1,5 %) | 275 (83,8 %) | 3 (0,9 %) |
| Pituitary | 283 | 204 (72,1 %) | 9 (3,2 %) | 8 (2,8 %) | 24 (8,5 %) | 87 (30,7 %) | 138 (48,8 %) | 17 (6 %) | 13 (4,6 %) | 21 (7,4 %) |
| Small Intestine Terminal Ileum | 187 | 120 (64,2 %) | 28 (15 %) | 22 (11,8 %) | 35 (18,7 %) | 56 (29,9 %) | 42 (22,5 %) | 4 (2,1 %) | 174 (93 %) | 2 (1,1 %) |
| Stomach | 359 | 227 (63,2 %) | 44 (12,3 %) | 39 (10,9 %) | 64 (17,8 %) | 128 (35,7 %) | 79 (22 %) | 5 (1,4 %) | 301 (83,8 %) | 3 (0,8 %) |
| Thyroid | 653 | 434 (66,5 %) | 47 (7,2 %) | 51 (7,8 %) | 110 (16,8 %) | 211 (32,3 %) | 212 (32,5 %) | 22 (3,4 %) | 358 (54,8 %) | 24 (3,7 %) |
| Whole Blood | 755 | 501 (66,4 %) | 68 (9 %) | 68 (9 %) | 113 (15 %) | 234 (31 %) | 249 (33 %) | 23 (3 %) | 412 (54,6 %) | 29 (3,8 %) |

*Data characteristics of the investigated tissues classified by age and separately by cause of death. Cause of death based on 4-point Hardy-Scale: 0) Cases on ventilator before death 1) violent and fast death, i.e. accident 2) fast death of natural causes 3) intermediate death with terminal phase of 1-24 hours 4) slow death after long illness.*

**Figure 1.**
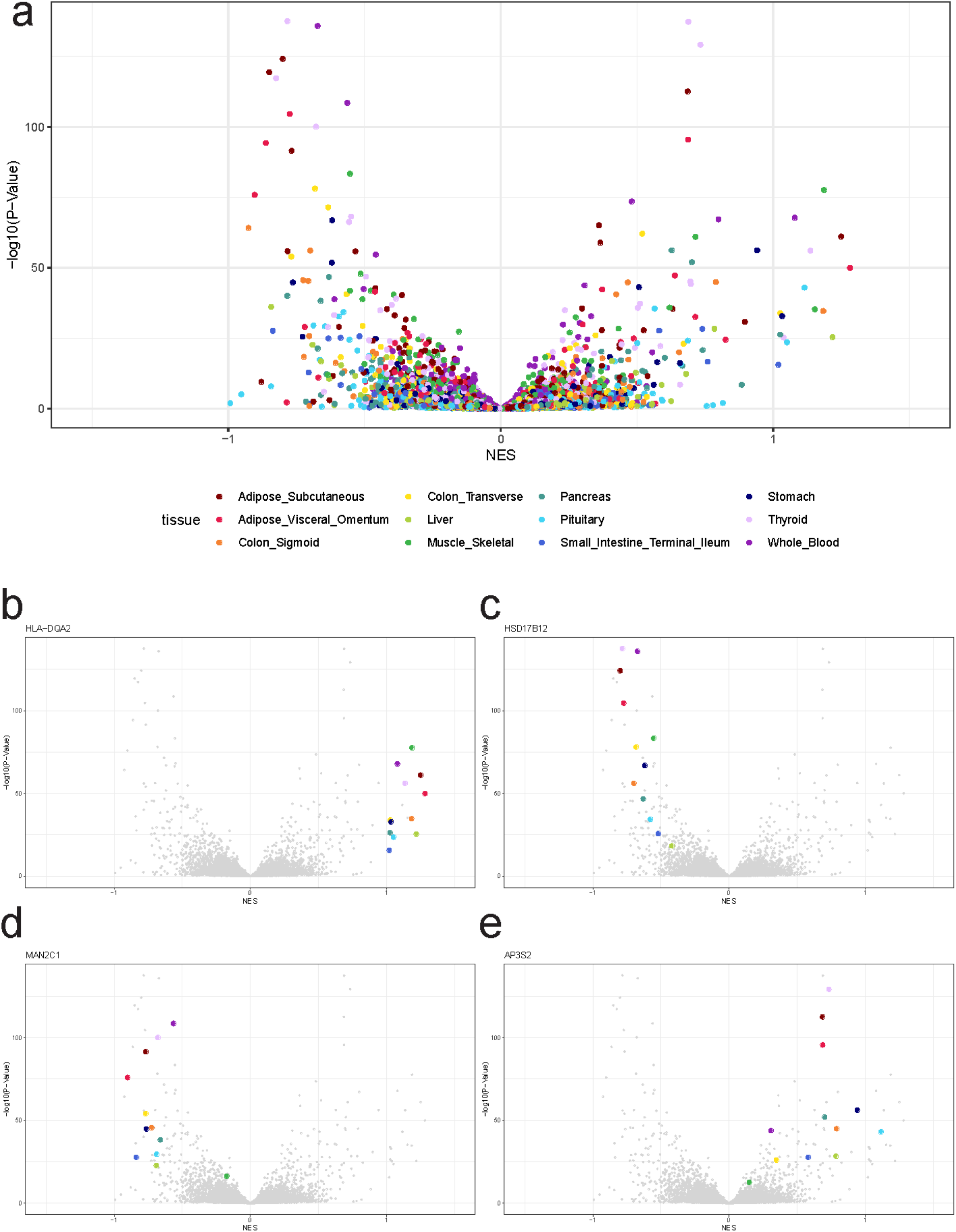
Volcano plot of all the 6664 calculated eQTL - eGene pairs, with the strongest tissue-wide eQTL NESs highlighted. **a)** Global overview **b)** rs601945 **-** *HLA-DQA2.* **c)** rs1061810 - *HSD17B12.***d)** rs13737 -*MAN2C1.* **e)** rs4932265 - *AP3S2.*

Four SNPs had strong (NESs) present in all tissues (**Fig. 1**). The strongest positive NESs were observed with a single SNP across all tissues between rs601945 and *HLA-DQA2*. The mean NES was 1.12 (SD=0.10, P ≤ 2.32·10^−16^, **Fig. 1b**). *HLA-DQA2* is involved in multiple disease- and immune response-related pathways [11]. Among the strongest negative NESs was rs1061810 which is an eQTL for *HSD17B12.*This eQTL - eGene pair had a mean NES of −0.64 (SD=0.11, P≤ 4.86·10^−19^, **Fig. 1c**) across all tissues. *HSD17B12* is involved in synthesis of fatty acids [11]. The strongest NES for this eQTL was observed in subcutaneous fat (NES = −0.80, P = 6.77·10^−125^). A strong eQTL NES was also observed between *MAN2C1* and rs13737 in all tissues, the mean NES was −0.68 (SD = 0.18, P≤ 6.49·10^−17^, **Fig. 1d**). The strongest normalized effect size of this pair was observed in subcutaneous fat (NES = −0.90, P =1.22·10^−76^), *MAN2C1* is involved in glycan degradation [11]. Lastly, *AP3S2* was observed to be influenced by the risk allele of rs4932265, the mean effect size was 0.65 (SD = 0.27, P≤ 3.02·10^−13^, **Fig. 1e**). *AP3S2* is thought to play a role in the lysosome [11].

### Type 2 diabetes-associated eQTLs link to metabolism and the ribosome pathway

In all tissues the majority of the SNPs mapped to *metabolic pathways*, with in absolute terms the highest numbers in whole blood (38 SNPs, **Fig. 2a)**, transverse colon (28 SNPs), thyroid (23 SNPs), stomach (22 SNPs) and pancreas (15 SNPs). Rs576123 (synonym for rs505922) was mapped in six tissues (transverse colon, pancreas, stomach, thyroid, whole blood and the Ileum) to metabolic pathways involving the *ABO* gene. Of note, a SNP in LD with rs576123 (rs8176719, R^2^ = 0.93, **Fig. 2b**) is a frameshift variant for *ABO.* The NES for ABO was positive (NES ≥ 0.17), except for whole blood (NES = −0.35, P = 6.98·10^−11^, **Fig. 2c**). The strongest NES for ABO was observed in the pancreas (NES = 0.74, P = 1.50·10^−21^, **Fig. 2d**).

**Figure 2.**
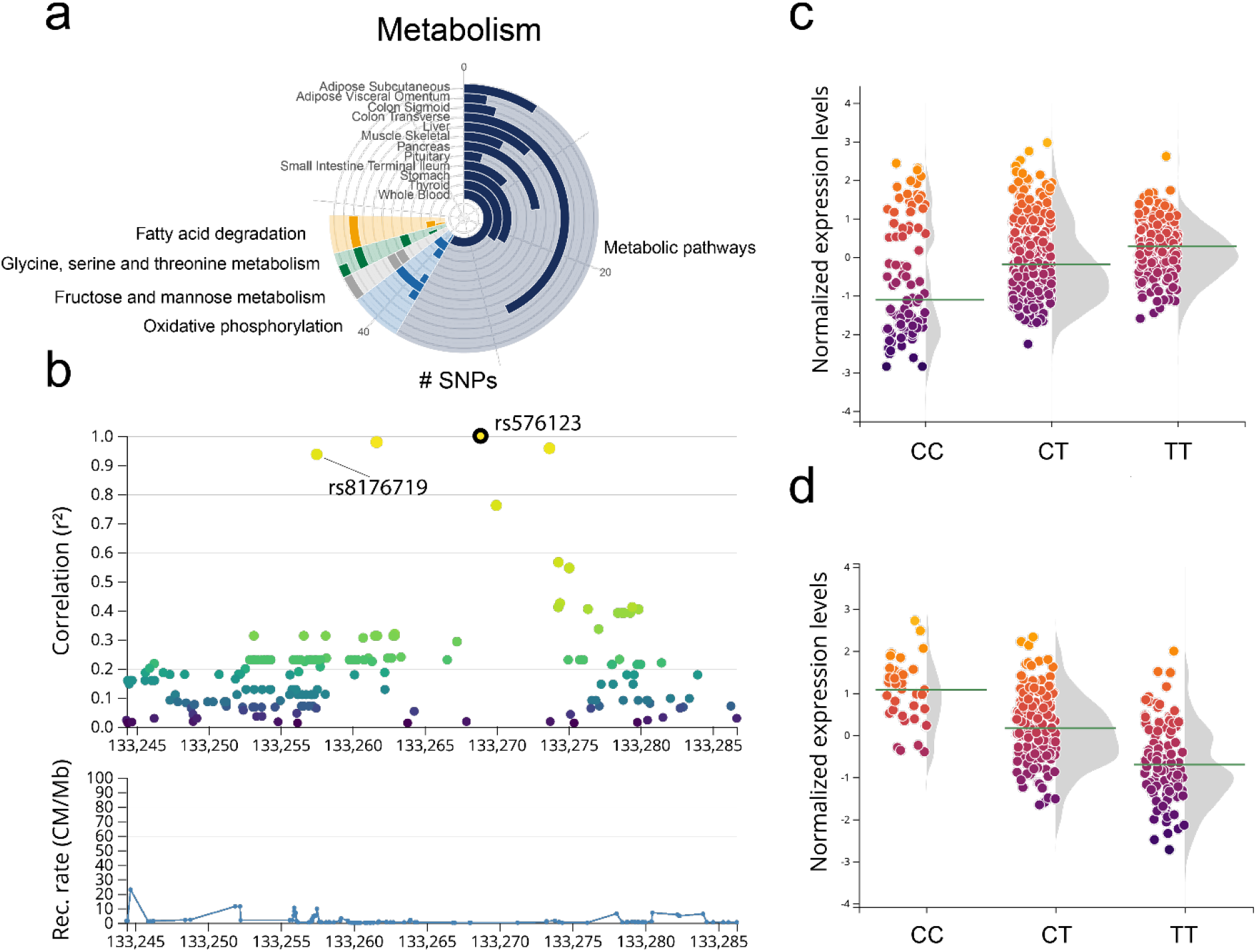
Overview of eQTLs mapped to metabolism and in-depth analysis of lead SNP rs576123. **a)** Overview of the number of eQTLs that are mapped to the corresponding pathways in the investigated tissues. **b)** Locus zoom of rs576123 with R^2^ of the surrounding SNPs and recombination rate, including rs8176719 (frameshift variant). **c)** Violin plot of the normalized expression levels of *ABO* in whole blood with the haplotypes of rs576123. **d)** Violin plot of the normalized expression levels of *ABO* in the pancreas with the haplotypes of rs576123.

The second most enriched process was *genetic information processing*. The ribosome pathway was enriched in eleven tissues (**Fig. 3a**). The modules in the various tissues that were enriched for the ribosome pathway all had varying numbers of associated eQTLs (n = 1-5) and eGenes (**Fig. 3b, Fig. 3c**). However, all modules shared a common eQTL - eGene pair, namely, rs12719778 and *RPL8.* The highest NES of rs12719778 on *RPL8* was observed in whole blood (NES = −0.10, P = 8.46·10^−13^, **Fig. 3d**). In nine of the eleven tissues in which the ribosome pathway was enriched, the modules also contained rs12920022 and *RPL13*, which had the strongest NES in skeletal muscle (NES = −0.26, P = 3.36·10^−26^, **Fig. 3e**).

**Figure 3.**
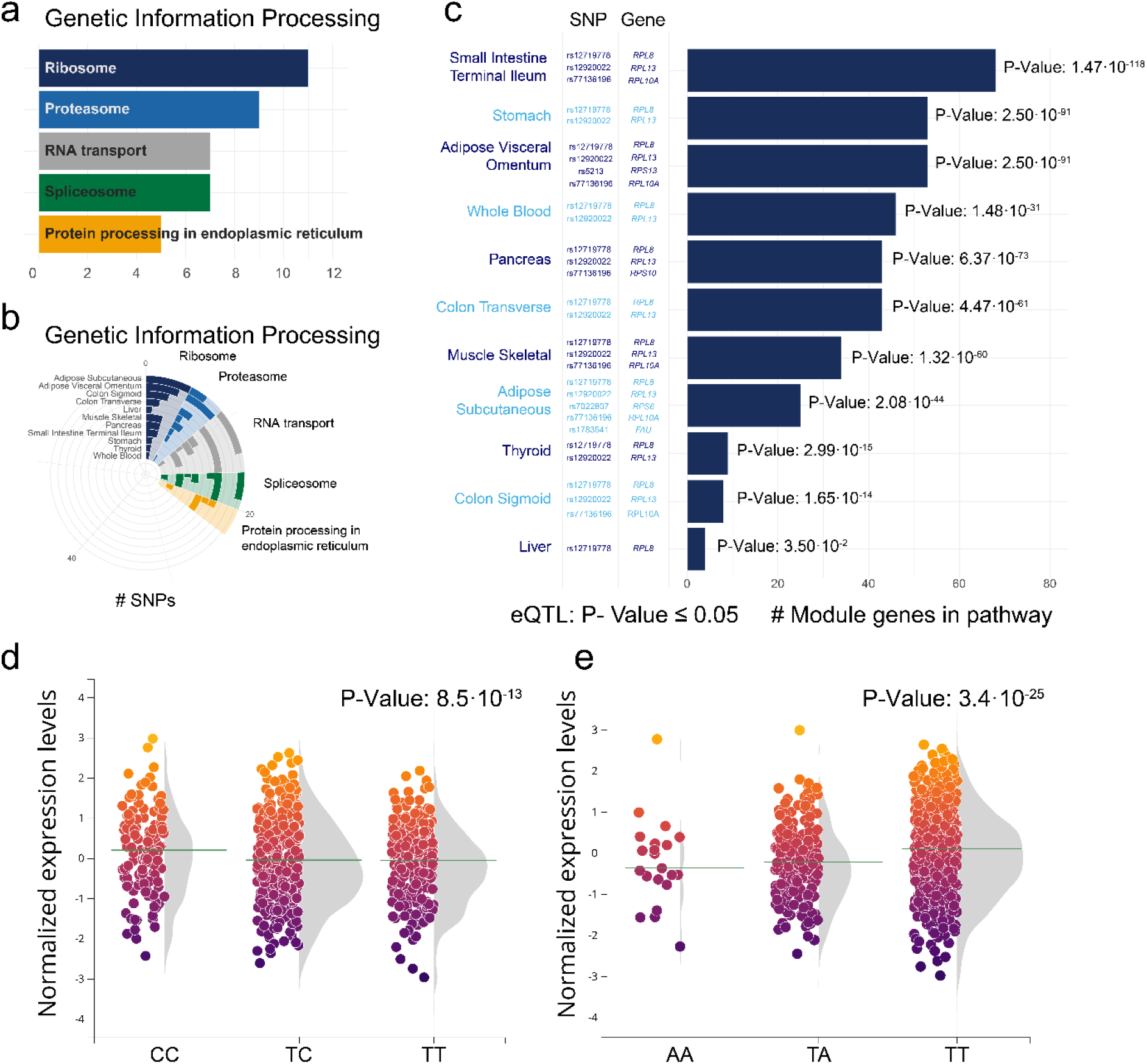
Overview of eQTLs mapped to Ribosome **a)** Number of tissues in which the corresponding pathways were enriched **b)** Overview of the number of eQTLs that are mapped to the corresponding pathways in the investigated tissues. **c)** Pathway enrichment results for ribosomes, with the eQTLs and eGenes involved. **d)** Violin plot of the normalized expression levels of *RPL8* in whole blood with the haplotypes of rs12719778. **e)** Violin plot of the normalized expression levels of *RPL13* in skeletal muscle with the haplotypes of rs12920022.

### In-depth analysis of rs601945 shows an association with primarily HLA genes

Among the enriched pathways, multiple pathways were immune-related (i.e. *Th 17 cell differentiation, Th 1 and Th 2 cell differentiation)*. All immune-related pathways that were enriched were linked with rs601945 and *HLA-DQA2.* As such, we explored the observed effect of rs601945 on the HLA gene *HLA-DQA2* in more detail (**Fig. 1b**). Rs601945 is located in an intergenic region and is in LD (R^2^ ≥ 0.8) with 94 SNPs. Four genes are located in this LD region: *HLA-DRB5, HLA-DRB1, HLA-DQA1* and *RNU1-61P.* For this LD region, 57 chromatin interactions are known in blood cells (CD34^+^, CD4^+^ memory, CD4^+^ naïve and CD4^+^ T-cells, **Fig. 4a**), 40 of which are interactions with loci located in HLA genes (*HLA-DQA1* = 19, *HLA-DQB1* = 13, *HLA-DRB1* = 6, *HLA-DQA2* = 3, *HLA-DRA* = 3, *HLA-DRB5* = 3, *HLA-DOB* = 1). In CD4^+^ memory cells 13 SNPs were located in enhancer regions and 3 SNPs in flanking active transcription start sites (TSS). In CD4^+^ naïve cells 11 SNPs were located in enhancer regions and 3 SNPs in flanking active TSSs. Sixty-six eGenes were identified that were influenced by rs601945 (P ≤ 0.05, cis = 25, trans = 41). Taking into account the more stringent adjusted P-value defined by GTEx, 19 eGenes remained (cis = 11, trans = 8, **Fig. 4b**). Among the significantly affected eGenes were primarily HLA genes. Rs601945 had positive and negative NESs with multiple *HLA* genes (**Fig. 4c**). As previously described, the strongest positive NESs were observed with *HLA-DQA2* with the strongest association observed in skeletal muscle (NES = 1.19, P = 2.50·10^−78^, **Fig. 4d**). The strongest opposite effect was observed with *HLA-DQB1* (NES = −0.50, P = 1.18·10^−16^, **Fig. 4e**). As rs601945 influences primarily HLA genes that are involved in many biological pathways, rs601945 was linked to seven pathways in multiple tissues. That is *cell adhesion molecules pathway* (**Fig. 4f**) in all twelve tissues, to five pathways, including *phagosome* and various immune pathways in eleven tissues (**Fig. 4g, Fig. 4h**) and linked to one pathway, *intestinal immune network for IgA production* in ten tissues (**Fig. 4h**).

**Figure 4.**
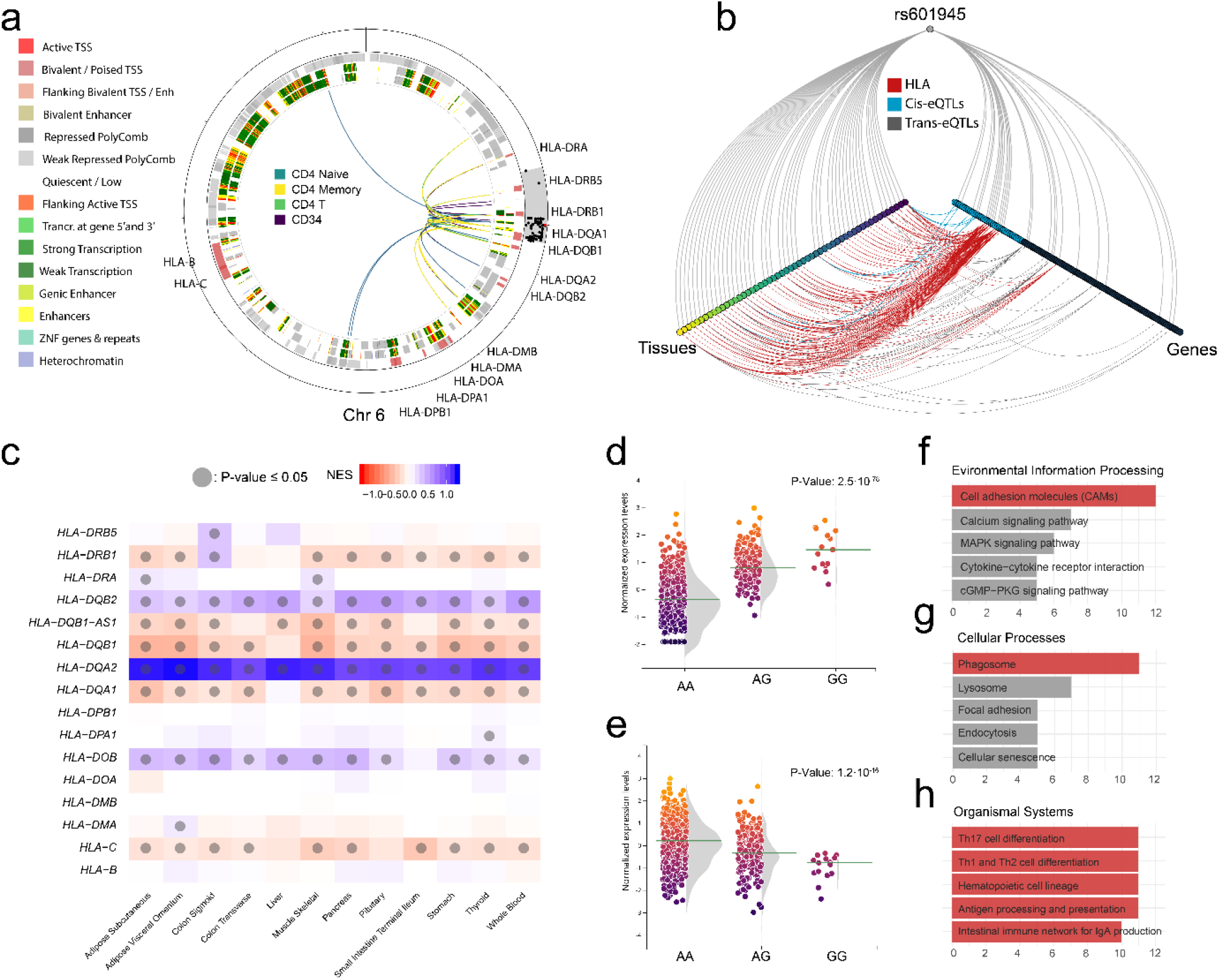
In-depth analysis of rs601945 **a)** Chromatin interactions in blood cells (CD34^+^, CD4^+^ memory, CD4^+^ naïve and CD4^+^ T) of the region that is in LD (R^2^ ≥ 0.8) with rs601945 and chromatin states of CD4^+^ naïve (outer-ring) and In CD4^+^ memory cells (inner-ring). **b)** Hive plot showing the association of rs601945 (center axis) with eGenes (right axis) in various tissues (left axis). **c)** NESs of all HLA genes investigated with rs601945. **d)** Violin plot of the normalized expression levels of *HLA-DQA2* in skeletal muscle with the haplotypes of rs601945. **e)** Violin plot of the normalized expression levels of *HLA-DQB1* in skeletal muscle with the haplotypes of rs12719778. **f, g, h)** Number of tissues in which the corresponding pathways were enriched with HLA associated pathways highlighted.

## DISCUSSION

In this study, we developed an R-package that aided us in understanding the functional consequences of T2D-associated SNPs. The R-package, called CONQUER collects up-to-date data, directed by SNPs of interest from a multitude of databases and repositories and analyses and visualizes the data. In contrast to previous studies that had similar approaches[12, 13], we have developed open-source software that is available as an R-package where only the SNPs and tissues of interest have to be specified and that can be used with minimal programming experience.

With CONQUER we developed a tool that is universal and versatile as it can be used for various diseases and phenotypes where SNPs are of interest. Because we included data from multiple sources of various molecular levels, it provides researchers with a broad range of information that aids them in understanding their phenotype of interest. Additionally, CONQUER expands the search space of consequential effects by including co-expressed genes of eGenes which might reveal up- and downstream consequences. With increasing amounts of data, the complexity also increases. To maintain clear overview of the data, we implemented two separate views 1) where in-depth analyses of single SNPs can performed and 2) where multiple SNPs and their aggregated consequences can be investigated and linked to biological pathways. CONQUER is dependent on the availability of the Application Programming Interfaces (API) to access databases (GTEx, Ensembl and LDlink). This is a strength, as the latest versions of these databases will always be accessed without changing the programming structure of CONQUER. However, if API access itself is changed or the databases are discontinued, then, updates to CONQUER are required. In contrast, access to the static data sources (e.g. meQTLs, miQTLs, pQTLs, chromatin states and chromatin interactions) is more secure as we maintain the source package (*conquer.db*). In addition, we will regularly update *conquer.db* as new studies will become available. All eQTLs are calculated with the GTEx API. Within GTEx sample sizes vary substantially per tissues, as a consequence, the number of significant eQTLs is correlated with the number of samples (R^2^=0.91). In the current study we take this into account by applying a liberal threshold and by assessing the NES across tissues. However, specific signatures of tissues with low sample counts might go unnoticed.

CONQUER was used to investigate 403 diabetes-associated SNPs in more detail. T2D is a metabolic disorder, in accordance, most SNPs linked to metabolic pathways. The metabolic pathways as curated by KEGG [11] consists of 1489 genes and is an encompassing term for all pathways that are involved in metabolism. Our results show that SNPs that are directly linked to metabolism do not influence a single metabolic process but are scattered among various metabolic pathways (e.g. oxidative phosphorylation, fatty acid degradation, fructose and mannose metabolism and glycine serine and threonine metabolism). Due to this dispersion of SNPs between numerous pathways it remains difficult to assign groups of SNPs to specific processes in specific tissues. This together with the variety of pathways to which SNPs are mapped shows that T2D has a lot of different points of engagement through which it can originate and progress, which is accordance with heterogeneous nature of T2D [14]. We also linked SNPs to different pathways classified as genetic information processing. As such, proteasome, RNA transport, spliceosome and protein processing in endoplasmic reticulum were pathways to which various SNPs were mapped. Additionally, seven SNPs were mapped to the ribosome pathway. The link between T2D SNPs and the ribosome pathway was observed in eleven tissues. Seven ribosomal genes with predominately negative effect sizes were associated with seven T2D GWAS hits. Although the association between ribosomal content and T2D has extensively been studied[15–17], genetic susceptibility to T2D has previously not been linked to a decreased expression of ribosomal genes. Moreover, the hormone insulin and ribosomal content are tightly connected. Insulin stimulates the synthesis of ribosomal proteins in various tissues[18, 19] and a loss of ribosomal proteins is associated with an inhibition of AKT phosphorylation activity and the insulin pathway[20]. Rs601945 was highlighted as it influences many HLA genes that are involved in multiple pathways. Rs601945 was associated with the HLA region. The HLA region has previously been associated with T2D[3, 21], however, our results reveal that the effects are wide-spread as its association with altered expression of various HLA genes was observed in all investigated tissues. Interestingly, while the HLA region represents the highest risk for T1D [22], our results are pointing to a connection between HLA-DQA2 and T2D. In addition, our pQTL analyses also highlighted immune response pathways. Our data support that T2D has an immunometabolic component involving, like T1D, members of both innate and adaptive immune response. Altogether, CONQUER revealed three biological main processes that could explain, in part, the association between SNPs and T2D susceptibility. In addition our results show that T2D SNPs influence metabolism through various pathways, that the ribosome pathway is influenced in multiple tissues through different combinations of SNPs and that rs601945 has wide-spread effects as it influences many genes that are involved in multiple immune related pathways. CONQUER was also used to analyze single SNP effects. Both *AP3S2* and *HSD17B12* have previously been found in relation to T2D, but in limited number of tissues. AP3S2 in human pancreatic islets[31] and HSD17B12 in adipose, liver and muscle tissue and, whole blood[3], which are relevant for the treatment of T2D [32]. However, the genetic consequences are not limited to these tissues as our results show. As such, rs1061810 was found to be associated with altered expression of *HSD17B12*. The effect of rs1061810 on *HSD17B12* has previously been described in adipose, liver and muscle tissue and, whole blood [3]. However, our results showed that the influence of rs1061810 on *HSD17B12* is not only present in these tissues but in all twelve tissues that we investigated. rs11037579 had lower expression of *HSD17B12* in all twelve tissues that we investigated, including adipose tissue. This result corroborates the finding that *HSD17B12* expression is downregulated in the adipose tissue of insulin-resistant subjects [23] and plays a role in adipogenesis [24]. The *HSD17B12* gene codes a bifunctional enzyme involved in the biosynthesis of estradiol and the elongation of very long chain fatty acids. One of the strongest observed effects was between rs4932265 and *AP3S2*. *AP3S2* is a subunit of the AP-3 complex which is involved in budding of vesicles from the Golgi membrane [25]. *AP3S2* has been linked to T2D in six different GWASs [21, 26–30] with various populations (South Asian, Japanese and European ancestry) and with four different SNPs, three of which (rs12912009, rs2028299, rs8031576) are in LD (R^2^ ≥ 0.80) with rs4932265. In the current study we established that *AP3S2* has a higher expression in the twelve tissues in individuals carrying the risk allele of rs4932265. Despite increasing evidence for the role *AP3S2* in T2D susceptibility it remains unclear how *AP3S2* is involved, although there is some evidence pointing at a beta-cell defect (wood et al https://doi.org/10.2337/db16-1452). In twelve tissues we have observed a negative effect size for *MAN2C1* with the T2D risk allele of rs13737. Downregulation of *MAN2C1* is known to cause delay in cell growth and inducing apoptosis [31, 32]. *MAN2C1* binds with *PTEN* and thereby inhibits its lipid phosphatase activity [31]. *PTEN* inhibits activation of PI3K-AKT signaling pathway[31, 33], a pathway known to be involved in T2D development[33]. As we observe a negative effect size for *MAN2C1,* it is suggested that in the twelve investigated tissues, for individuals carrying the risk allele of rs13737 the PI3K-AKT signaling pathway could be inhibited in part, by a reduced expression of *MAN2C1* through *PTEN.* This could explain the association of rs13737 with T2D susceptibility. In six tissues we linked rs576123, located in intronic region of the ABO gene to metabolic pathways. While, *ABO* is at the basis of the ABO blood group system as it indirectly encodes for blood group antigens [34], a recent study has observed an impaired insulin secretion within O blood type subjects. In this study, SNPs located within the first intron have been connected to a reduced activity of the glycosyltransferases encoded by the ABO gene and specific targeting of the ABO gene by shRNA has led to a reduced glucose stimulated insulin secretion[35]. The effect of reduced ABO expression in the other tissues needs to be established.

## CONCLUSION

The R-package CONQUER allows efficient integration of multiple datasets. With data on various levels, visualized in a tidy manner, we were able to uncover potential consequences of T2D associated risk loci. As such, SNPs could be linked through various biological mechanisms to insulin resistance and insulin secretion and comprehend the increased T2D risk. Our findings highlight the importance of an integrative approach where risk loci for T2D are not only seen as individual risk factors but also as a network of risk factors. With CONQUER we developed software that does this, uses the latest available data and is easy to use.

## METHODS

### CONQUER

*CONQUER* was developed in R version 3.6.1 and is available from Git (https://github.com/roderickslieker/CONQUER). The user end the package consists of two intuitive function calls, *summarize* and *visualize*. The summarize function minimally requires a list of SNPs (rs* IDs), a directory to store them in and a token from LDlink to allow access to their API. Additional options *multiAnalyze* (boolean), to allow integrated analysis of multiple SNPs. This option, requires a list of tissues in which the integrated analysis should be performed. Summarize will collect all data described below for each SNP and store this in a small file that can be used in a later stage. The visualize function invokes a Shiny-based dashboard, with interactive plots of the integrated analysis (if performed) and a tab where individual SNPs can be visualized. Interactive figures were made using JavaScript Data-Driven Documents (d3.js) version 4.13.0, based on existing and newly developed plots. D3.js code was integrated in R making use of the htmlwidgets R-package[36] and all tools were integrated into the R package *CONQUER.d3*. Interactive heatmaps were made using plotly[37]. The interactive circos plot was made with the R-package BioCircos[38]. Interactive tables were generated with the DT package[39].

### Data acquisition

The data acquired for *CONQUER* are based on the human genome reference build GRCh38/hg38. The data is both collected from static sources and Application Programming Interfaces (APIs). The static sources are available in a separate R data package called *conquer.db*. *CONQUER* loads this data package when needed. As *conquer.db* is a separate package it is easily updated with the latest datasets without altering the programming structure of CONQUER. Static data include chromatin interactions, chromatin state segmentations, expression data, transcription factor binding sites, pQTLs, miQTLs. The chromatin interactions were obtained from the 4D genome database[7]. To have data from multiple tissues (N=31), only IM-PET data was included in CONQUER. Originally this data was based on the human genome reference build GRCh19/hg19. UCSC LiftOver tool[40] was used to lift over the data to GRCh38/hg38. Chromatin state segmentations were obtained from the Roadmap Epigenomics Project for all cell types available (N=127, 15-state model) [6]. Normalized (TPM, Transcript per Million) expression data of all available tissues (N=54) was obtained from GTEx v8[5]. Missing expression values were imputed with k-nearest neighbor and default parameters of the *impute.knn* function from the R-package impute[41]. Data of pQTLs [44–47], meQTLs[48], miQTLs [49–52] were acquired from their corresponding references.

The remaining data (linkage disequilibrium, gene information, eQTLs) are obtained from APIs and are collected once the user has given the command. Elementary information about the SNP of interest is acquired from the Ensembl API [8]. The linkage disequilibrium (LD) structure originates from the LDlink API[51]. For both the Ensembl API and LDlink API the population can be specified, by default the population is set on Utah Residents with Northern and Western European Ancestry (CEU) from the 1000 Genomes Project phase 3[52]. The eQTLs and eGenes corresponding to the SNP of interest are computed making use of GTEx API. By default, GTEx has an eQTL mapping window of one Mb upstream and downstream of the transcription start site of a gene. In *CONQUER*, we expanded the search space by including genes that have chromosomal interaction with the LD region (R^2^ ≥ 0.80) of the leading SNP. *CONQUER* automatically sends a request to the GTEx API to calculate eQTLs for every available tissue utilizing GTEx v8[5]. Lastly, phenotype associations are acquired from the GWAS-catalog[53]. For every queried SNP, *CONQUER* generates an *RData* object containing all previously described data and stores it in a directory the user has provided.

### Statistical analyses

First, CONQUER was used to retrieve all the available data of the 403 risk loci associated with T2D [1]. Next, the modularization and pathway enrichment were performed by CONQUER on twelve T2D relevant tissues (**Table 1**). It should be noted that GTEx reports normalized effect sizes (NES) as effect of the alternative allele relative to the reference allele. However, a variant associated with T2D can be either the reference allele or alternative allele. Therefore, we investigated the effect size of the risk allele of T2D relative to the other allele based on an additive model. Also, due to the variability of sample sizes of GTEx across tissues and their association with P-values of eQTLs, we use both normalized effect size and P-values for the interpretation of the results. Figures were directly from CONQUER or additionally made using ggplot2.

### Modularization and pathway enrichment

With multiple SNPs, *CONQUER* can modularize SNPs and associate them with biological pathways in tissues of interest (**Fig. 5**). All tissues in GTEX can be included. Based on the GTEx data, eQTLs and their associating eGenes are selected (P-value ≤ 0.05). For these eGenes, co-expressed genes are identified by performing correlation analyses with imputed GTEx expression data in the corresponding tissues. Co-expression between genes is assumed when *rho* ≥ *0.90.* Next, the eGenes and their co-expressed genes are hierarchical clustered[54, 55] based the distance between genes (*1 – rho*). The number of modules within the clustered data is optimized by maximizing the globalSEmax of the gap statistic[56] using the *cluster R package[57]*. Modules of co-expressed genes and eGenes are then tested for pathway enrichment based on KEGG pathways. For each pathway odds ratios and accompanying P-values are calculated with Fisher’s exact test[58]. If a module does not contain an eQTL or is not enriched for a pathway, it is omitted from the analysis.

**Figure 5.**
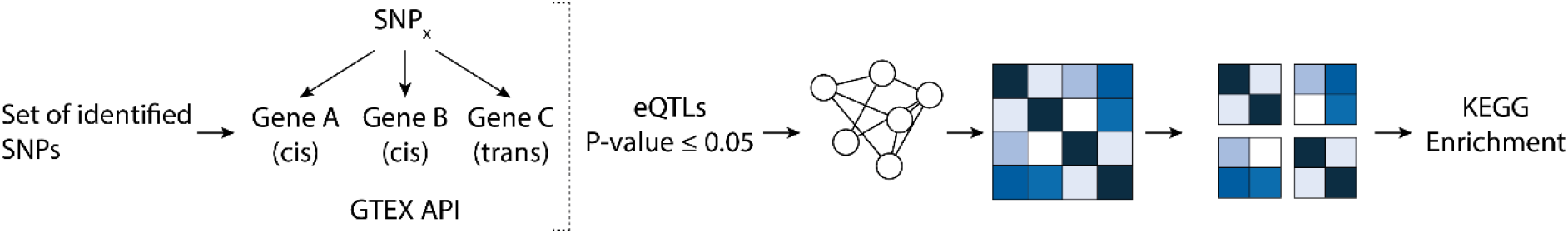
Schematic overview of SNP modularization. Starting with a set of SNPs, cis- and trans genes are calculated with the GTEx API. After applying a threshold (P ≤ 0.05), sets of co-expressed genes are determined after which all included genes (eGenes and co-expressed genes) are clustered. Each module (cluster) resulting from the cluster analysis is then enriched with KEGG.

## ACKNOWLEDGEMENTS

This project has received funding from the Innovative Medicines Initiative 2 Joint Undertaking under grant agreement No 115881 (RHAPSODY). This Joint Undertaking receives support from the European Union’s Horizon 2020 research and innovation programme and EFPIA. This work is supported by the Swiss State Secretariat for Education‚ Research and Innovation (SERI) under contract number 16.0097-2. The opinions expressed and arguments employed herein do not necessarily reflect the official views of these funding bodies.

## AUTHOR CONTRIBUTIONS

GAB, RCS, LMTH designed the study. GAB and RCS analyzed the data and wrote the draft manuscript. All authors critically read and revised the manuscript and approved the final version of the manuscript.

## REFERENCES

1. Mahajan A, Taliun D, Thurner M, Robertson NR, Torres JM, Rayner NW, Steinthorsdottir V, Scott RA, Grarup N, Cook JP, et al: Fine-mapping of an expanded set of type 2 diabetes loci to single-variant resolution using high-density imputation and islet-specific epigenome maps. Nat Genet 2018, 50:1505–1513.

2. Xue A, Wu Y, Zhu Z, Zhang F, Kemper KE, Zheng Z, Yengo L, Lloyd-Jones LR, Sidorenko J, Wu Y, et al: Genome-wide association analyses identify 143 risk variants and putative regulatory mechanisms for type 2 diabetes. Nature Communications 2018, 9.

3. Scott RA, Scott LJ, Mägi R, Marullo L, Gaulton KJ, Kaakinen M, Pervjakova N, Pers TH, Johnson AD, Eicher JD, et al: An Expanded Genome-Wide Association Study of Type 2 Diabetes in Europeans. Diabetes 2017, 66:2888–2902.

4. Maurano MT, Humbert R, Rynes E, Thurman RE, Haugen E, Wang H, Reynolds AP, Sandstrom R, Qu H, Brody J, et al: Systematic Localization of Common Disease-Associated Variation in Regulatory DNA. Science 2012, 337:1190–1195.

5. Consortium GT: The Genotype-Tissue Expression (GTEx) project. Nat Genet 2013, 45:580–585.

6. Bernstein BE, Stamatoyannopoulos JA, Costello JF, Ren B, Milosavljevic A, Meissner A, Kellis M, Marra MA, Beaudet AL, Ecker JR, et al: The NIH Roadmap Epigenomics Mapping Consortium. Nature Biotechnology 2010, 28:1045–1048.

7. Teng L, He B, Wang J, Tan K: 4DGenome: a comprehensive database of chromatin interactions. Bioinformatics 2015, 31:2560–2564.

8. Hunt SE, McLaren W, Gil L, Thormann A, Schuilenburg H, Sheppard D, Parton A, Armean IM, Trevanion SJ, Flicek P, Cunningham F: Ensembl variation resources. Database (Oxford) 2018, 2018.

9. Watanabe K, Taskesen E, van Bochoven A, Posthuma D: Functional mapping and annotation of genetic associations with FUMA. Nature Communications 2017, 8.

10. Lu HC, Herrera Braga J, Fraternali F: PinSnps: structural and functional analysis of SNPs in the context of protein interaction networks. Bioinformatics 2016, 32:2534–2536.

11. Kanehisa M, Goto S: KEGG: Kyoto Encyclopedia of Genes and Genomes. Nucleic Acids Research 2000, 28:27–30.

12. Fernandez-Tajes J, Gaulton KJ, van de Bunt M, Torres J, Thurner M, Mahajan A, Gloyn AL, Lage K, McCarthy MI: Developing a network view of type 2 diabetes risk pathways through integration of genetic, genomic and functional data. Genome Med 2019, 11:19.

13. Cirillo E, Kutmon M, Gonzalez Hernandez M, Hooimeijer T, Adriaens ME, Eijssen LMT, Parnell LD, Coort SL, Evelo CT: From SNPs to pathways: Biological interpretation of type 2 diabetes (T2DM) genome wide association study (GWAS) results. PLoS One 2018, 13:e0193515.

14. DeFronzo RA, Ferrannini E, Groop L, Henry RR, Herman WH, Holst JJ, Hu FB, Kahn CR, Raz I, Shulman GI, et al: Type 2 diabetes mellitus. Nature Reviews Disease Primers 2015, 1:15019.

15. Stirewalt WS, Wool IG, Cavicchi P: The relation of RNA and protein synthesis to the sedimentation of muscle ribosomes: effect of diabetes and insulin. Proceedings of the National Academy of Sciences of the United States of America 1967, 57:1885–1892.

16. Wool IG, Kurihara K: Determination of the number of active muscle ribosomes: effect of diabetes and insulin. Proceedings of the National Academy of Sciences of the United States of America 1967, 58:2401–2407.

17. Ozoe A, Sone M, Fukushima T, Kataoka N, Chida K, Asano T, Hakuno F, Takahashi S-I: Insulin Receptor Substrate-1 Associates with Small Nucleolar RNA Which Contributes to Ribosome Biogenesis. Frontiers in Endocrinology 2014, 5:24.

18. Proud CG: Regulation of protein synthesis by insulin. Biochemical Society transactions 2006, 34:213–216.

19. Proud CG, Denton RM: Molecular mechanisms for the control of translation by insulin. The Biochemical journal 1997, 328 (Pt 2):329–341.

20. Heijnen HF, van Wijk R, Pereboom TC, Goos YJ, Seinen CW, van Oirschot BA, van Dooren R, Gastou M, Giles RH, van Solinge W, et al: Ribosomal Protein Mutations Induce Autophagy through S6 Kinase Inhibition of the Insulin Pathway. PLOS Genetics 2014, 10:e1004371.

21. Zhao W, Rasheed A, Tikkanen E, Lee J-J, Butterworth AS, Howson JMM, Assimes TL, Chowdhury R, Orho-Melander M, Damrauer S, et al: Identification of new susceptibility loci for type 2 diabetes and shared etiological pathways with coronary heart disease. Nature Genetics 2017, 49:1450–1457.

22. Bradfield JP, Qu H-Q, Wang K, Zhang H, Sleiman PM, Kim CE, Mentch FD, Qiu H, Glessner JT, Thomas KA, et al: A Genome-Wide Meta-Analysis of Six Type 1 Diabetes Cohorts Identifies Multiple Associated Loci. PLOS Genetics 2011, 7:e1002293.

23. Elbein SC, Kern PA, Rasouli N, Yao-Borengasser A, Sharma NK, Das SK: Global gene expression profiles of subcutaneous adipose and muscle from glucose-tolerant, insulin-sensitive, and insulin-resistant individuals matched for BMI. Diabetes 2011, 60:1019–1029.

24. Söhle J, Machuy N, Smailbegovic E, Holtzmann U, Grönniger E, Wenck H, Stäb F, Winnefeld M: Identification of new genes involved in human adipogenesis and fat storage. PloS one 2012, 7:e31193–e31193.

25. Odorizzi G, Cowles CR, Emr SD: The AP-3 complex: a coat of many colours. Trends in Cell Biology 1998, 8:282–288.

26. Kooner JS, Saleheen D, Sim X, Sehmi J, Zhang W, Frossard P, Been LF, Chia K-S, Dimas AS, Hassanali N, et al: Genome-wide association study in individuals of South Asian ancestry identifies six new type 2 diabetes susceptibility loci. Nature Genetics 2011, 43:984–989.

27. Hara K, Fujita H, Johnson TA, Yamauchi T, Yasuda K, Horikoshi M, Peng C, Hu C, Ma RCW, Imamura M, et al: Genome-wide association study identifies three novel loci for type 2 diabetes. Human Molecular Genetics 2013, 23:239–246.

28. Bonàs-Guarch S, Guindo-Martínez M, Miguel-Escalada I, Grarup N, Sebastian D, Rodriguez-Fos E, Sánchez F, Planas-Fèlix M, Cortes-Sánchez P, González S, et al: Re-analysis of public genetic data reveals a rare X-chromosomal variant associated with type 2 diabetes. Nature Communications 2018, 9:321.

29. Suzuki K, Akiyama M, Ishigaki K, Kanai M, Hosoe J, Shojima N, Hozawa A, Kadota A, Kuriki K, Naito M, et al: Identification of 28 new susceptibility loci for type 2 diabetes in the Japanese population. Nature Genetics 2019, 51:379–386.

30. Kichaev G, Bhatia G, Loh P-R, Gazal S, Burch K, Freund MK, Schoech A, Pasaniuc B, Price AL: Leveraging Polygenic Functional Enrichment to Improve GWAS Power. The American Journal of Human Genetics 2019, 104:65–75.

31. Hopkins BD, Hodakoski C, Barrows D, Mense SM, Parsons RE: PTEN function: the long and the short of it. Trends in biochemical sciences 2014, 39:183–190.

32. Wang L, Suzuki T: Dual functions for cytosolic α-mannosidase (Man2C1): its down-regulation causes mitochondria-dependent apoptosis independently of its α-mannosidase activity. The Journal of biological chemistry 2013, 288:11887–11896.

33. Huang X, Liu G, Guo J, Su Z: The PI3K/AKT pathway in obesity and type 2 diabetes. International journal of biological sciences 2018, 14:1483–1496.

34. Patenaude SI, Seto NOL, Borisova SN, Szpacenko A, Marcus SL, Palcic MM, Evans SV: The structural basis for specificity in human ABO(H) blood group biosynthesis. Nature Structural Biology 2002, 9:685–690.

35. Li-Gao R, Carlotti F, de Mutsert R, van Hylckama Vlieg A, de Koning EJP, Jukema JW, Rosendaal FR, Willems van Dijk K, Mook-Kanamori DO: Genome-Wide Association Study on the Early-Phase Insulin Response to a Liquid Mixed Meal: Results From the NEO Study. Diabetes 2019, 68:2327.

36. Vaidyanathan R, Xie Y, Allaire J, Cheng J, Russell K: htmlwidgets: HTML Widgets for R. https://CRANR-projectorg/package=htmlwidgets 2018.

37. Sievert C: plotly for R.https://plotly-rcom 2018.

38. Cui Y, Chen X, Luo H, Fan Z, Luo J, He S, Yue H, Zhang P, Chen R: BioCircos.js: an interactive Circos JavaScript library for biological data visualization on web applications. Bioinformatics 2016, 32:1740–1742.

39. Xie Y, Cheng J, Tan X: DT: A Wrapper of the JavaScript Library ‘DataTables’.2019.

40. Kuhn RM, Haussler D, Kent WJ: The UCSC genome browser and associated tools. Brief Bioinform 2013, 14:144–161.

41. Hastie T TR, Narasimhan B, Chu G: impute: impute: Imputation for microarray data R package version 1.60.0. Bioconductor 2019.

42. An integrated encyclopedia of DNA elements in the human genome. Nature 2012, 489:57–74.

43. Yao C, Chen G, Song C, Keefe J, Mendelson M, Huan T, Sun BB, Laser A, Maranville JC, Wu H, et al: Genome-wide mapping of plasma protein QTLs identifies putatively causal genes and pathways for cardiovascular disease. Nat Commun 2018, 9:3268.

44. Wu L, Candille SI, Choi Y, Xie D, Jiang L, Li-Pook-Than J, Tang H, Snyder M: Variation and genetic control of protein abundance in humans. Nature 2013, 499:79–82.

45. Suhre K, Arnold M, Bhagwat AM, Cotton RJ, Engelke R, Raffler J, Sarwath H, Thareja G, Wahl A, DeLisle RK, et al: Connecting genetic risk to disease end points through the human blood plasma proteome. Nat Commun 2017, 8:14357.

46. Carayol J, Chabert C, Di Cara A, Armenise C, Lefebvre G, Langin D, Viguerie N, Metairon S, Saris WHM, Astrup A, et al: Protein quantitative trait locus study in obesity during weight-loss identifies a leptin regulator. Nat Commun 2017, 8:2084.

47. Huan T, Rong J, Liu C, Zhang X, Tanriverdi K, Joehanes R, Chen BH, Murabito JM, Yao C, Courchesne P, et al: Genome-wide identification of microRNA expression quantitative trait loci. Nat Commun 2015, 6:6601.

48. Borel C, Deutsch S, Letourneau A, Migliavacca E, Montgomery SB, Dimas AS, Vejnar CE, Attar H, Gagnebin M, Gehrig C, et al: Identification of cis-and trans-regulatory variation modulating microRNA expression levels in human fibroblasts. Genome Res 2011, 21:68–73.

49. Gamazon ER, Ziliak D, Im HK, LaCroix B, Park DS, Cox NJ, Huang RS: Genetic architecture of microRNA expression: implications for the transcriptome and complex traits. Am J Hum Genet 2012, 90:1046–1063.

50. Liu Cea: MirSNP, a database of polymorphisms altering miRNA target sites, identifies miRNA-related SNPs in GWAS SNPs and eQTLs. BMC genomics 2012.

51. Machiela MJ, Chanock SJ: LDlink: a web-based application for exploring population-specific haplotype structure and linking correlated alleles of possible functional variants. Bioinformatics 2015, 31:3555–3557.

52. Clarke L, Fairley S, Zheng-Bradley X, Streeter I, Perry E, Lowy E, Tasse AM, Flicek P: The international Genome sample resource (IGSR): A worldwide collection of genome variation incorporating the 1000 Genomes Project data. Nucleic Acids Res 2017, 45:D854–D859.

53. Buniello A, MacArthur JA L, Cerezo M, Harris LW, Hayhurst J, Malangone C, McMahon A, Morales J, Mountjoy E, Sollis E, et al: The NHGRI-EBI GWAS Catalog of published genome-wide association studies, targeted arrays and summary statistics 2019. Nucleic Acids Research 2019, 47:D1005–D1012.

54. Leonard Kaufman PJR: Finding Groups in Data: An Introduction to Cluster Analysis. New York: John Wiley & Sons, Inc.; 1990.

55. Belbin L, Faith DP, Milligan GW: A comparison of two approaches to beta-flexible clustering. Multivariate Behavioral Research 1992, 27:417–433.

56. Dudoit S, Fridlyand J: A prediction-based resampling method for estimating the number of clusters in a dataset. Genome Biology 2002, 3:research0036.0031.

57. Maechler M: Cluster R-Package. Cran 2019.

58. Fisher RA: The Logic of Inductive Inference. Journal of the Royal Statistical Society 1935, 98:39–82.

